# Alternative splicing of NUMB correlates with tumor immune evasion and metabolic adaptation

**DOI:** 10.1101/2025.08.11.669729

**Authors:** Yangjing Zhang, C. Jane McGlade

## Abstract

Alternative splicing of the *NUMB* gene at exon 9 generates protein isoforms linked to tumor growth and metastasis, but its impact on the tumor immune microenvironment remains unclear. Using RNA sequencing data from over 5,000 tumors across 16 cancer types, including breast cancer subtypes, we found that tumors with high *NUMB* exon 9 inclusion exhibit lower expression of immune-related genes, reduced immune cell infiltration, and decreased cytolytic activity, consistent with an immune “cold” phenotype. This pattern was consistent across multiple cancer types and breast cancer subtypes. Additionally, exon 9-high tumors showed evidence of increased oxidative phosphorylation, suggesting metabolic adaptations that support tumor progression. These findings identify *NUMB* exon 9 inclusion as a potential biomarker of immune evasion and highlight opportunities for patient stratification and targeted therapies based on *NUMB* exon 9 inclusion levels.

## Introduction

Immune checkpoint inhibitors have revolutionized cancer treatment, offering new therapeutic avenues for malignancies previously considered untreatable. Despite their transformative impact, only a subset of patients experience a meaningful clinical benefit from immune checkpoint blockade therapy^1^. The ability to predict which patients will respond to immunotherapy remains a significant challenge, as multiple factors influence treatment outcomes. Tumors classified as immune “hot” are characterized by high levels of immune cell infiltration, particularly T cells, which makes them more likely to respond favorably to immunotherapies^1^. Additionally, the presence of an active immune environment enhances the effectiveness of conventional treatments and clinical trials have shown that combining chemotherapy with immunotherapy improves patient outcomes, particularly in cancers such as triple-negative breast cancer ^2,3^ and non-small cell lung cancer ^4^. This synergistic interaction highlights the crucial role of the tumor’s immune landscape in guiding effective treatment strategies.

The *NUMB* gene plays a crucial role in cell fate determination and undergoes alternative splicing (AS) of two exon cassettes, coding exon 3 and exon 9, resulting in the production of four distinct mRNA and protein isoforms. During development, splicing regulation of *NUMB* exon 9 leads to a reduction in exon 9–containing isoform expression in differentiated tissues ^5^. However, the re-expression of *NUMB* exon 9 has been identified as a frequent recurrent AS event in cancer ^5–7^. A pan-cancer analysis of exon 9 inclusion across 27 cancer types from The Cancer Genome Atlas (TCGA) revealed a significant increase in *NUMB* exon 9 inclusion in at least 15 cancer types compared to normal tissues ^7,8^. Additionally, hazard ratio (HR) analysis suggests that elevated exon 9 inclusion correlates with a modest but significant pan-cancer survival risk ^8^. In breast cancer specifically, higher exon 9 inclusion correlates with worse clinical outcomes consistent with the increased metastatic potential of *NUMB* exon 9 expressing cells in a xenograft human breast cancer mouse model. Mechanistically, several studies suggest that exon 9–included isoforms may lack the lysosomal targeting function observed in exon 9–skipped isoforms, leading to an increased surface presentation of oncogenic receptors and membrane proteins, which may promote tumor aggressiveness^7,9^. However, the tumorigenic effects of *NUMB* isoforms in an immunocompetent model have not yet been explored.

The present study investigates the relationship between *NUMB* exon 9 inclusion and tumor immunity. Using RNA sequencing data from a variety of human tumors in the TCGA dataset, we examined the differentially expressed genes associated with *NUMB* exon 9 inclusion. Our analysis reveals that tumors with high exon 9 inclusion exhibit reduced expression of genes associated with immune cell infiltration and immune cytolytic activity across 11 cancer types. This trend was also observed in breast cancer subtypes, where increased exon 9 inclusion correlates with a less active immune environment. These findings suggest a novel mechanism by which *NUMB* inclusion may contribute to tumour progression, linking alternative splicing of exon 9 to tumor immune modulation. Our study highlights the potential of *NUMB* exon 9 inclusion as a biomarker for patient stratification and underscores its potential importance as a target for precision therapies aimed at improving immune responses and treatment outcomes in diverse cancer types.

## Results

### Differential gene expression in *NUMB* exon9 high vs low tumor groups

The Percent Spliced In (PSI) index represents the proportion of a gene’s mRNA transcripts that include a specific exon, providing a quantitative measure of alternative splicing events ^7^. To investigate the genes and pathways that are differentially activated in tumors with different *NUMB* exon9 inclusion level, we analyzed publicly available RNA sequencing data for 5491 patient tumors across 16 cancer types ^8,10^. Within each cancer type, patients were stratified into two subgroups based on *NUMB* exon 9 PSI values: those in the top 10th percentile were classified as exon 9-high, while those in the bottom 10th percentile were classified as exon 9-low (***Figure 1A***). The exon9 high vs low samples within each cancer type were separated by at least 25% in PSI value, except in GBM which was separated by only 5% and therefore used as a negative control ^8^. To identify differentially expressed genes (DEGs), we utilized the DESeq2 R package to conduct quantification and statistical inference of systematic changes between exon9 high vs exon9 low groups, as compared to within-group variability. For each cancer type, a list of genes that were significantly differentially expressed (false discovery rate/FDR < 0.1) was generated. These gene lists were used in a pathway enrichment analysis to infer pathways that are differentially activated in each condition ^11^ (***Figure 1A***).

**Figure 1.**
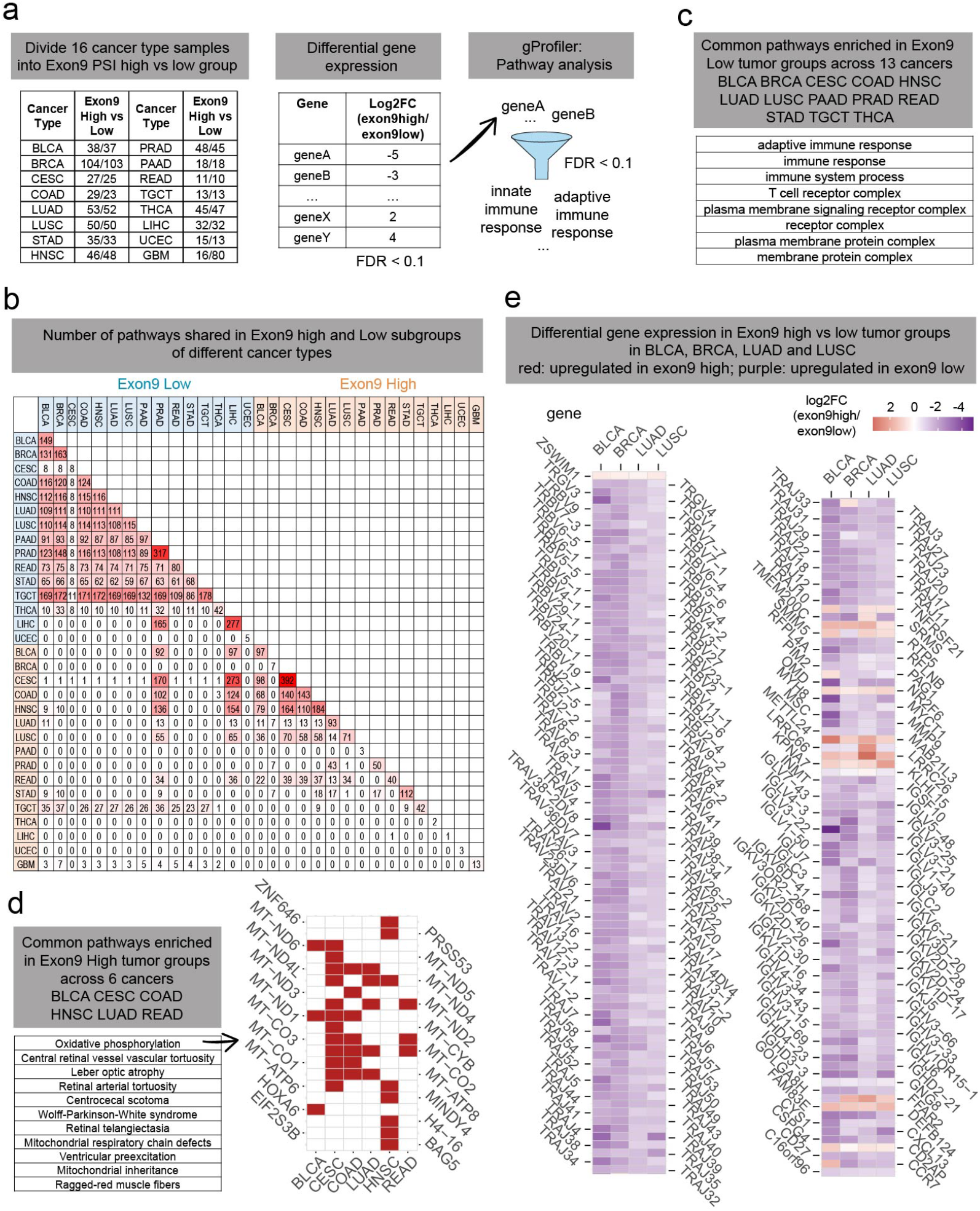
Differentially expressed genes and pathways in ex9 high vs low tumor groups in different cancer types. **(A)** Workflow for the analysis of differential pathways in exon9 high versus low subgroups among 16 TCGA tumor types. The number of samples in exon9 high vs low subgroups were tabulated within each tumor type. Exon9 PSI high group includes samples with top 10 percentile of exon9 inclusion level, whereas exon9 PSI low group represents samples with bottom 10 percentile of exon9 inclusion level within specific cancer type. DESeq2 R packaged was used to quantify and infer statistical changes between the two conditions as compared to within-condition variability. The output is a list of genes for each cancer type that are differentially expressed in exon9 high vs low subgroups with log2 fold change values and false discovery rate (FDR) value. Only genes with FDR < 0.1 are considered significant and used for pathway analysis. We use gProfiler to perform pathway enrichment analysis to our ranked gene lists. **(B)** Number of common pathways shared in exon9 high vs low subgroups of different cancer types. Cancer types in blue are upregulated pathways in exon9 low subgroup. Cancer types in orange are upregulated pathways in exon9 high subgroup. Shades of red represent the number of pathways: 0, white; biggest number, red. **(C)** Common pathways upregulated in exon9 low tumor groups across 13 TCGA tumor types. **(D)** Common pathways upregulated in exon9 high tumor groups across 6 TCGA tumor types. Right. Genes involved in oxidative phosphorylation are upregulated in exon9 high tumor group. Red, identified to be upregulated in exon9 high subgroup within certain cancer types. **(E)** Genes shared in BLCA, BRCA, LUAD and LUSC tumor types that are differentially expressed between exon9 high vs low subgroups within each tumor type. The log2FC (exon9high/exon9low) was represented as a heatmap. BLCA bladder urothelial carcinoma; BRAC breast invasive carcinoma; CESC cervical squamous cell carcinoma and endocervical adenocarcinoma; COAD colon adenocarcinoma; HNSC head and neck squamous cell carcinoma; LIHC liver hepatocellular carcinoma; LUAD lung adenocarcinoma; LUSC lung squamous cell carcinoma; PAAD pancreatic adenocarcinoma; PRAD prostate adenocarcinoma; READ rectum adenocarcinoma; STAD stomach adenocarcinoma; UCEC uterine corpus endometrial carcinoma; THCA thyroid carcinoma; TGCT testicular germ cell tumors; GBM glioblastoma multiforme;

Out of the 16 cancer types analyzed, multiple pathways were identified as upregulated in the exon9 low group (***Figure 1B***). Strikingly, 13 out of the 16 cancer types analyzed exhibited significant enrichment in adaptive immune response and immune response pathways in the exon9 low subgroup, except for LIHC, UCEC and GBM (***Figure 1C)***. The specific genes differentially enriched in exon9 low group of tumors include T cell receptor (TCR) subunits and immunoglobin subunits, key signatures of immune T cell and B cells, suggesting a more immunologically active tumor microenvironment in the exon9 low subgroup.

Conversely, tumors with higher exon 9 inclusion exhibited upregulation of distinct metabolic pathways in some of the cancer types. One of the commonly enriched pathways across BRCA, CESC, COAD, HNSC, LUAD and READ cancer types was oxidative phosphorylation pathway, which included mitochondrial genes. This suggests that tumors with higher NUMB exon 9 inclusion may adapt to increased energy demands by upregulating mitochondrial metabolism (***Figure 1D)***, potentially promoting tumor aggressiveness and survival.

To further explore shared transcriptional changes across different cancers, we examined BLCA, BRCA, LUAD, and LUSC, which exhibited common DEG patterns associated with *NUMB* exon 9 inclusion levels (***Figure 1E***). In addition to T and B cell receptor genes, immune response related genes such as CD27 and CXCL13 were more highly expressed in exon9 low subgroups (***shown in purple***), while genes involved in signaling pathways unrelated to immune activation, such as SRMS (Src-Related Kinase Lacking C-Terminal Regulatory Tyrosine And N-Terminal Myristylation Sites), MAB21L3 (Mab-21 Like 3) and CD2AP (Cas Ligand With Multiple SH3 Domains) are more highly expressed in exon9 high subgroups (***shown in red***).

### *NUMB* Exon 9 Inclusion Correlates with Reduced Immune Cell Infiltration and Cytolytic Activity

Bulk tumor RNA sequencing data can also identify which immune cell types are differentially represented in tumors with different exon9 inclusion levels. Rooney *et al* consolidated marker genes for specific immune cell types as those with expression at least 2-fold greater than observed in any other cell type, and calculated enrichment scores (represented as a z-score) of a cell type meta-gene in a given TCGA sample using single sample gene set enrichment analysis (ssGSEA) ^12^. We integrated the enrichment score of various immune cell types for each tumor sample with the *NUMB* exon9 inclusion DEG data to compare the differential enrichment of immune cell types in exon9 high vs low subgroups within each cancer type. The average enrichment score of immune cell types in the exon9 high vs low subgroups within each cancer was determined (***Figure A)***. Within BLCA, BRCA, COAD, LUAD, LUSC, STAD, CESC, READ, SKCM and HNSC tumors, markers of most immune cell types are less enriched in tumors from exon9 high subgroup compared to exon9 low subgroup. This suggests a general absence of immune cell infiltration in tumors with high *NUMB* exon 9 inclusion.

Next, we hypothesized that exon9 expression could be a predictor of outcome in spontaneous tumor immunity. To test this, we utilized the cytolytic (cell lysis) activity score (CYT) calculated by Rooney et al., which quantifies immune cytotoxicity based on the gene expression levels of two key cytolytic effectors, granzyme A (GZMA) and perforin (PRF1). The expression of both cytolytic molecules is upregulated upon CD8+ T cell activation and during productive clinical responses to anti-CTLA-4 and anti-PD-L1 immunotherapies and correlates with improved prognosis ^12^. In 11 out of 19 tumor types analyzed, tumors with high exon9 inclusion level have a markedly lower immune cytolytic activity than those with low exon9 inclusion, including BLCA, KIRC, COAD, BRCA, LUAD, HNSC, STAD, LUSC, SKCM, THCA and PRAD (***Figure B)***.

Taken together, these findings suggest that the association between high *NUMB* exon9 inclusion and reduced immune cell infiltration is linked to a corresponding decrease in immune cytolytic activity.

### Less immune cell infiltration in breast cancer subgroups with high exon9 inclusion

Increased exon9 inclusion is a consistent feature across all five major breast cancer subtypes - basal like, HER2 enriched, luminal A, luminal B, and normal like - and is associated with significantly worse progression free survival in patients ^3^. To better understand the biological consequences of elevated exon 9 inclusion, we next investigated whether tumors with high exon 9 inclusion display distinct gene expression profiles and pathway activity compared to those with low exon 9 inclusion across breast cancer subtypes. Patient samples within each breast cancer subtype were stratified into exon 9-high and exon 9-low groups and differential gene expression analysis between groups were performed (***Figure 3A***). For each subtype, we identified genes with significant expression differences (FDR < 0.1) between exon 9 high vs low subgroups and ranked them by fold change for pathway enrichment analysis.

**Figure 2.**
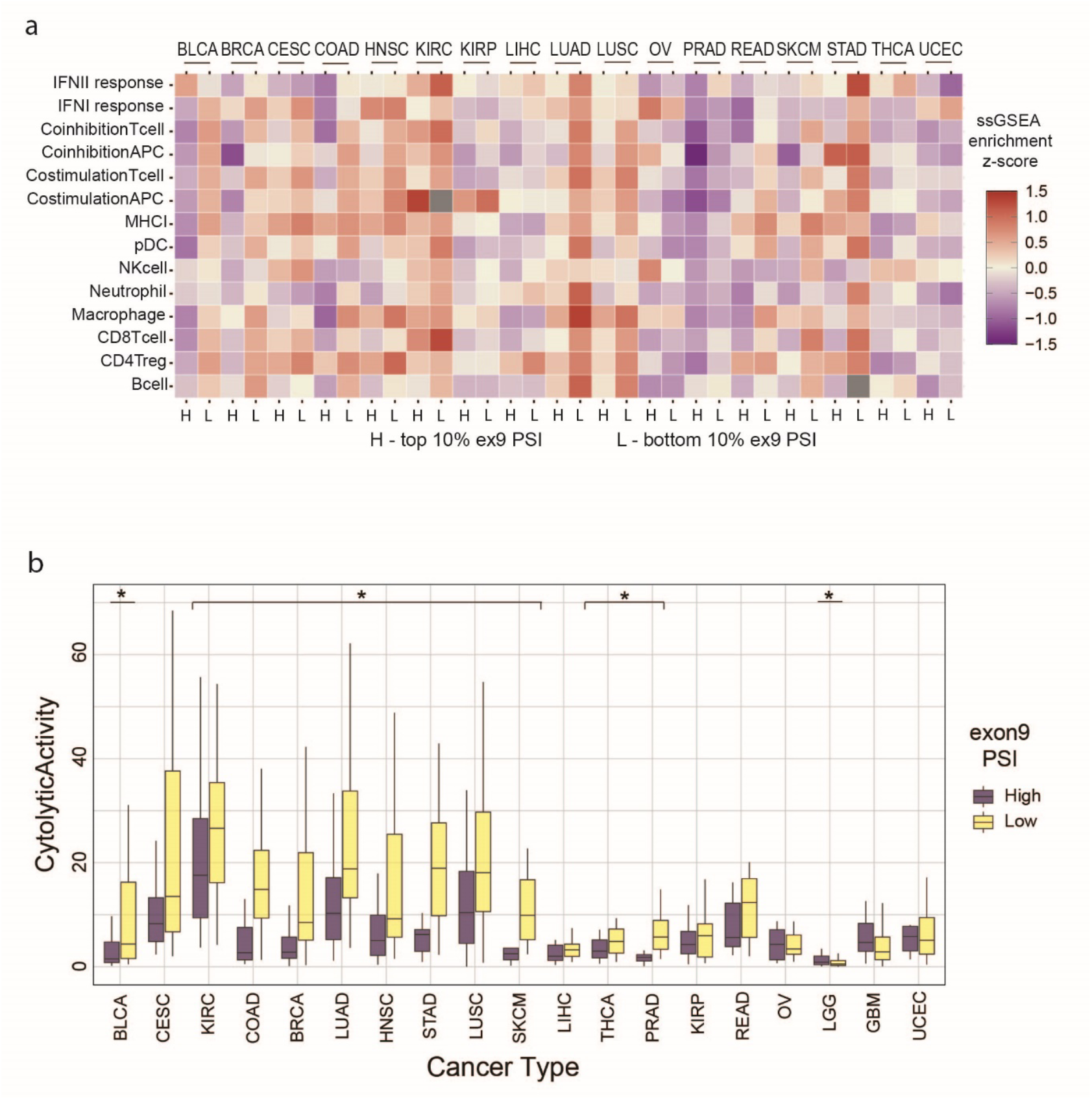
Reduced immune cell enrichment and cytolytic activity correlates with high *NUMB* exon9 expression. **(A)** Relationship between exon9 inclusion and immune cell type enrichment. Mean ssGSEA enrichment z-score is plotted as heatmap between exon9 high vs low groups across 17 TCGA tumor types. Grey, no data. **(B)** Relationship between exon9 inclusion level and immune cytolytic activity. Cytolytic activity for exon9 high vs low groups across 19 TCGA tumor types. * Wilcoxon test p value < 0.05. KIRC kidney renal clear cell carcinoma; KIRP kidney renal papillary cell carcinoma; LGG brain lower grade glioma; OV ovarian serous cystadenocarcinoma; SKCM skin cutaneous melanoma

**Figure 3.**
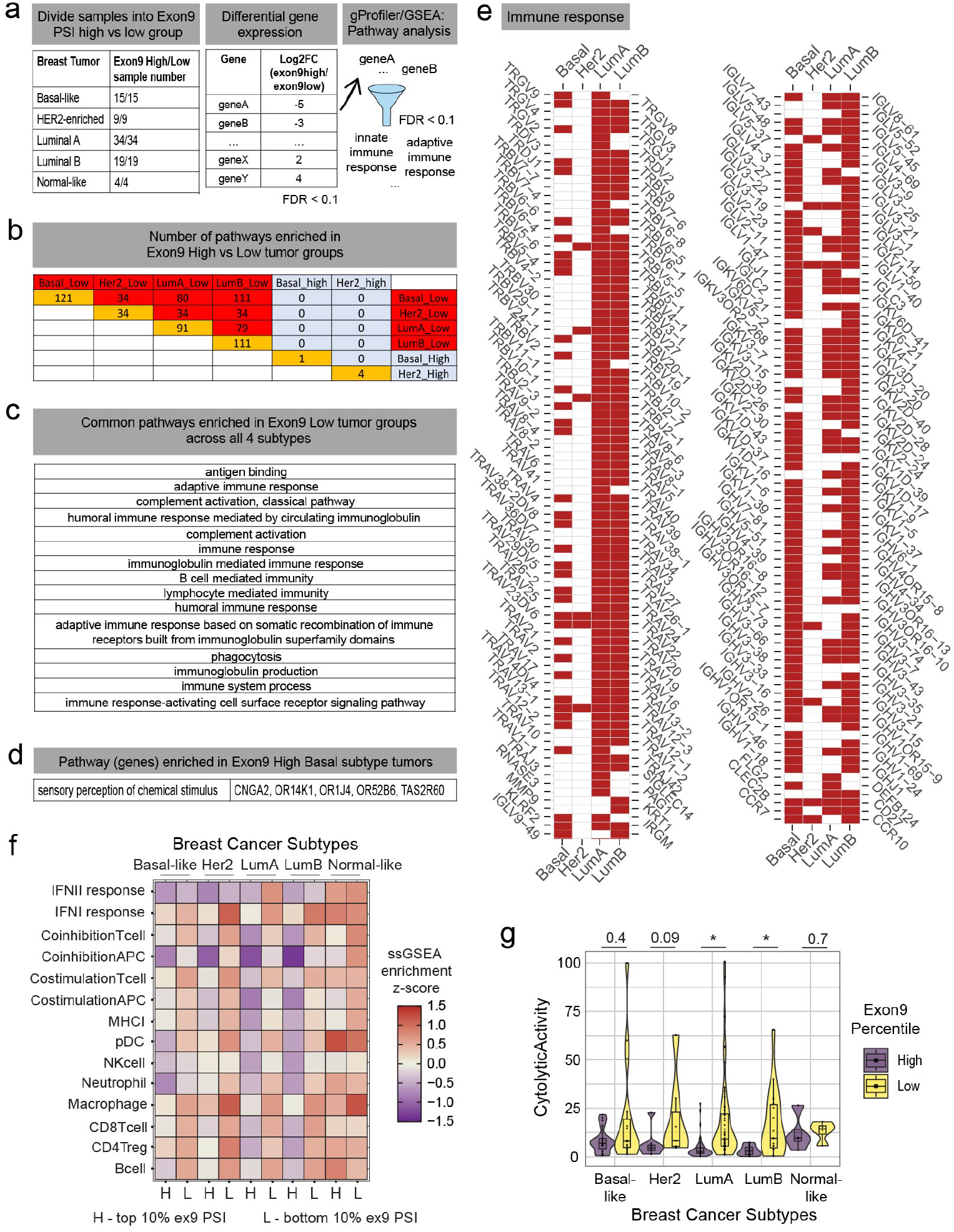
Less immune cell infiltration in breast tumors with higher *NUMB* exon9 inclusion level. **(A)** Workflow for analysis of differential pathways activated in exon9 high versus low subgroups in breast cancer subtypes. **(B)** Number of pathways enriched in exon9 high versus low tumor subgroups. **(C)** Common pathways upregulated in exon9 low tumor groups across four breast cancer subtypes. **(D)** Pathway enriched in exon9 high basal like breast cancer patient. Genes identified in this pathway are listed. **(E)** Immune response genes upregulated in exon9 low groups within each breast cancer subtypes. Red indicates genes identified in that subgroup. **(F)** Relationship between exon9 inclusions and immune cell type enrichment. Mean ssGSEA enrichment z-score is plotted as heatmap between exon9 high vs low groups in each breast cancer subtype. **(G)** Immune cytolytic activity measured by transcript level of *GZMA* and *PRF1* between exon9 high vs low groups within each breast cancer subtype. *, t test p-value < 0.05.

Fewer differentially expressed genes were identified in the normal like breast cancer group likely due to both the smaller separation of exon9 PSI values between exon9 high and low subgroups and the limited sample size, leaving no pathways identified for this subtype. In addition, fewer pathways were upregulated in exon9 high groups compared to exon9 low groups across subtypes (***Figure B***). Interestingly, genes encoding taste and olfactory receptors were upregulated in exon9 high group of basal like breast cancer (***Figure D***), suggesting subtype-specific effects related to exon 9 inclusion beyond immune-related functions.

Consistent with our findings in breast cancer (BRCA) as a whole, the most commonly downregulated pathways in exon 9-high tumors (indicated as enriched pathways in exon 9 low subgroup) across all four major subtypes were strongly associated with immune responses (**Figure 3C**). Specifically, tumors with high exon 9 inclusion exhibited reduced expression of genes encoding T cell receptor subunits and immunoglobulin subunits, which serve as signatures for T cells and B cells, respectively (***Figure 3E***). To better define the immune cell landscape of exon 9-high and exon 9-low tumors, we integrated immune cell enrichment scores from Rooney et al ^12^ with exon 9 inclusion data and visualized the differences in immune cell populations across subtypes (***Figure 3F***). In all subtypes except the normal-like group, exon 9-low tumors demonstrated higher enrichment of multiple immune cell types, including T cells, B cells, macrophages, dendritic cells, costimulatory and coinhibitory T cell subsets, and interferon I (IFNI) response cells (***Figure 3G***). In line with these observations, immune cytolytic activity, as measured by transcript levels of GZMA and PRF1, was lower in tumors with high exon 9 inclusion across subtypes, though this difference did not reach statistical significance in the normal-like and basal-like subtypes (***Figure 3F***).

Together, these results suggest that breast tumors—particularly within the luminal subtypes—with high *NUMB* exon 9 inclusion are characterized by reduced immune cell infiltration and diminished immune cytolytic activity. These features point to an impaired anti-tumor immune response, potentially contributing to the worse clinical outcomes observed in these patients.

## Discussion

The correlation between reduced immune signatures and high NUMB exon 9 inclusion across at least 11 cancer types and four major breast cancer subtypes suggests that the Exon9 included *NUMB* isoforms may play an active role in promoting an immune-excluded tumor microenvironment. In these tumors, diminished infiltration of immune cells, particularly cytotoxic lymphocytes, is accompanied by reduced immune cytolytic activity, as reflected by lower expression of GZMA and PRF1. These findings point to a global suppression of anti-tumor immunity in tumors with elevated NUMB exon 9 inclusion, which may facilitate tumor progression through immune evasion, activation of compensatory non-immune pathways, and ultimately contribute to the poorer prognosis observed in these patients.

While previous work has demonstrated that *NUMB* exon 9 promotes tumor growth and metastasis in xenograft models of lung cancer and breast cancer, these models lack an intact immune system ^7,13^, limiting the ability to assess a role in regulating anti-tumor immunity. To determine whether *NUMB* exon 9 inclusion directly alters immune responses within tumors, future studies utilizing immunocompetent models will be required to investigate the causal relationship between *NUMB* splicing and immune cell exclusion within a functional immune system. Further, mass cytometry (CyTOF) and single-cell RNA sequencing could be employed to define the specific immune cell populations affected by *NUMB* exon 9 inclusion, clarifying whether specific immune subsets, such as CD8+ T cells, macrophages, or dendritic cells, are preferentially excluded or suppressed. Additionally, testing the efficacy of immune checkpoint inhibitors in *NUMB* exon 9-high versus exon 9-low tumors could provide critical insights into whether exon 9 inclusion status predicts responsiveness to immunotherapy.

Beyond immune evasion, prior studies have shown that NUMB exon 9-included isoforms enhance the surface localization of oncogenic membrane proteins, including adhesion molecules, transporters, and growth receptors ^7^. This raises the possibility that patients with high *NUMB* exon 9-expressing tumors may be particularly susceptible to antibody–drug conjugates ^14^ targeting surface markers, such as ITGβ5. In contrast, patients with low *NUMB* exon 9 inclusion and robust immune infiltration might derive greater benefit from immunotherapies, such as checkpoint blockade or adoptive T cell transfer. Moreover, our pathway analysis revealed that *NUMB* exon 9-high tumors exhibit increased oxidative phosphorylation activity, suggesting a metabolic vulnerability that could be exploited therapeutically. Given this metabolic shift, tumors with high exon 9 inclusion may be more sensitive to inhibitors targeting oxidative phosphorylation, opening opportunities for targeted metabolic interventions.

In summary, our findings position *NUMB* exon 9 inclusion as an important molecular feature linked to immune suppression, metabolic rewiring, and poor prognosis across multiple cancer types. These insights suggest that *NUMB* exon 9 splicing status could serve as a biomarker for patient stratification, guiding the selection of immunotherapies, and targeted therapies in precision oncology approaches. Future mechanistic studies in immunocompetent models and clinical validation will be essential to translate these findings into therapeutic benefit.

## Methods

### Data download

Comprehensive alternative splicing analysis was conducted across RNA sequencing data from The Cancer Genome Atlas and GTEx database ^10^. The PSI values for exon9 (chr14:73745988 - 73746132) and exon3 (chr14:73783097 - 73783130) regions for *NUMB* was extracted from the supplementary tables together with sample IDs. The raw read counts of all aligned genes were downloaded from the supplementary data portal. For BRCA, information for the molecular stratification of breast cancer samples into different subtypes was downloaded through TCGAbiolink R package.

With TCGA solid tumor patient samples, cytolytic activity (CYT) was calculated as the geometric mean of GZMA and PRF1 (as expressed in TPM, 0.01 offset) by Rooney et al ^12^. Marker genes for immune cell types were defined based on at least 2-fold higher expression compared to other cell types, using Fantom5 and DMAP datasets. Up to 10 marker genes per immune cell type were selected from Fantom5, and their enrichment in each sample was assessed using single-sample gene set enrichment analysis (ssGSEA), followed by z-score normalization across samples ^12^.

### Differential gene expression analysis

R package DESeq2 (version 1.24.0) was used for differential gene expression analysis between exon9 PSI high vs low groups in different breast tumor subtypes. Within each cancer type, patients are stratified into exon9 high (top 10 percentile) and exon9 low (bottom 10 percentile) groups based on exon9 PSI value of *NUMB* from the primary tumor. The transcript expression table (aligned but unnormalized readcounts) for TCGA patient samples were downloaded ^10^ and samples in the two exon9 inclusion groups were extracted. Differential expression comparison between exon9 high vs low groups within each cancer type was performed. Only the differentially expressed genes with FDR value of lower than 0.1 was used for later analysis. Two methods were used to analyze pathway enrichment using the ranked gene lists from DESeq2 analysis: G:profiler and GSEA with default settings ^11^. Similar analysis was done on different breast cancer subtypes.

### Immuno-profile analysis

By combining the published data on cytolytic activity and enrichment score for immune cell types ^12^ to the previous TCGA data frame with Numb exon9 inclusion information, the same tumor stratification into exon9 PSI high and low subgroups was performed. Comparison of cytolytic activity and immune cell type enrichment between the two groups across 19 tumor types was carried out using R. Within breast cancer, the comparison of cytolytic activity and immune cell type enrichment between the two groups across 5 subtypes was carried out.

## Acknowledgements

This work was supported by funding from Canadian Institutes of Health Research (Project Grant FRN 106507) and the Canadian Cancer Society (Challenge Grant - 707475) to CJM.

